# Ecological change and conflict reduction led to the evolution of a transformative social behavior in ants

**DOI:** 10.1101/2022.09.25.509371

**Authors:** Marie-Pierre Meurville, Daniele Silvestro, Adria C. LeBoeuf

## Abstract

Behavioral innovations can be ecologically transformative for lineages that perform them and for their associated communities. Many ecologically dominant, superorganismal, and speciose ant lineages use a mouth-to-mouth fluid exchange behavior – trophallaxis – to share both exogenously sourced and endogenously produced materials across their colonies, while lineages that are less abundant, less cooperative and less speciose tend not to perform this behavior. How and why this behavior evolved and fixed in only some ant lineages remains unclear and whether this trait enables ants’ ecological dominance is not yet understood. Here we show that trophallaxis evolved in two major events ~110 Ma in lineages that today encompass 36% of ants, and in numerous smaller and more recent events. We found that trophallaxis evolved early only in ant lineages that had reduced intra-colonial conflict by losing workers ability to reproduce. Our causal models indicate that this signature behavior of superorganismal ants required social cooperation and ecological opportunism, and likely contributed to the large colony sizes and speciation patterns of the ants that use it and dominate our landscapes today. We hypothesize that the early evolution of trophallaxis was brought about by a major shift in terrestrial ecosystems through the origin and diversification of flowering plants and the consequent opportunistic inclusion of nectar and sap-sucker honeydew in the ant diet.

## Introduction

Changes in intra- and inter-species behavior have had dramatically impacts on the ecology of our planet. Novel behaviors can impact not only a species’ own evolutionary trajectory through selection, speciation and extinction dynamics, but also the evolutionary trajectories of associated species, and even entire ecosystems^1–3^. The evolution from carnivorous wasps to nectar- and pollen-foraging bees^4^ was transformational for plant reproduction with the evolution and co-evolution of countless pollination syndromes and adaptations and likely promoted the diversification of many flowering plant clades^5,6^. Occasionally new behaviors and their associated morphology can be so crucial, that through selection, speciation and extinction dynamics, entire clades exclusively display central adaptations around the behavior. This occurred in the evolution of mammals when a novel social transfer of material evolved – milk^7–10^. Lactation and nursing behaviors were so essential to the mammalian lineage that these traits have been retained across the 300 Ma long evolutionary history and among the >4000 living species^7^.

Behavioral traits and associated phenotypic adaptations were central to the evolution of one of the most ecologically dominant invertebrate clades: the ants^3,11,12^. The ant phylogeny holds extreme diversity in ecology, morphology, life-history traits, but also in behavior. At one extreme are the living descendants of early-branching lineages with small, often monomorphic colonies of predatory specialists, and at the other extreme are the more derived ecosystem-dominating generalist species with colossal colonies and high levels of queen-worker dimorphism. The latter show dramatic reproductive division of labor, to the point where a single colony can be considered a superorganism, where the sexuals are the germline and sterile workers are the soma^13^.

A majority of ecosystem-dominating generalist ants perform a behavior called stomodeal trophallaxis, wherein adult workers regurgitate the contents of their crop to nestmates in a mouth-to-mouth interaction^14^, though it is not clear why these traits appear to co-vary. It has been suggested that trophallaxis behavior evolved to transport sugary liquids acquired through mutualisms with sapsuckers or with plants^15,16^. Species that use trophallaxis tend not to have a sting and tend to have sterile workers or workers that cannot be mated, suggesting that these traits might prohibit the evolution of trophallaxis^14^. In some ants where trophallactic fluid has been characterized (*Camponotus*), it has been shown to contain not only food but also endogenously produced materials including proteins, small molecules, RNA and even hormones that shift with colony maturation^17–19^. This suggests that in such superorganismal species, trophallaxis behavior may enable a form of social circulatory system or shared metabolism across the colony^14,19^, and large colonies have been proposed to require more coordination^20–22^. At present, the evolutionary dynamics that led to the single or repeated origins of this behavioral trait remain largely unexplored. Similarly unclear is the link between trophallaxis and the emergence of superorganismality. Finally, it is not known whether trophallaxis, like lactation and nursing in mammals, is an irreversible key innovation or a volatile trait in ants.

In this study, we tested whether trophallaxis behavior enabled the shift from predaceous ant ancestors to more modern generalists, assessed which pressures led trophallaxis to evolve and how this behavior impacted speciation. We combine deep learning predictions, phylogenetic comparative methods and causal models to infer the trophallaxis behavior across hundreds of ant species and reconstruct its evolution and impact on the long diversification history of this clade.

## Results

After compiling all records of trophallaxis from the literature (265 papers, 205 species, Supp Table1), we used a Bayesian neural network to train a predictive model able to infer the presence or absence of this behavior based on phylogenetic, ecological, and morphological data and life-history traits, which could be collected for a larger number of species. These included the inclusion of sugary liquid food in the diet, colony size, presence of workers with full reproductive potential, presence of sting, and proxies for climate preferences (see Methods and Supplementary Discussion for more information).

Our predictive model reached a cross-validation test accuracy of 84% (Supp Table 2). Since a BNN inherently quantifies the uncertainty around predictions, we could identify a posterior probability threshold above which the expected accuracy increases to 95%. Using this threshold, 24% of the species in the test set were classified as uncertain. To further assess the accuracy of our model we reached out to myrmecologists with expertise on certain species groups to have their determination of trophallaxis behavior and compared their assessment with our predictions. Our model’s myrmecologist accuracy was 88% and included both false negatives and false positives (Supp Table 2)

Using feature permutation, we identified phylogeny (Δaccuracy of 4.5%), the inclusion of sugary liquid in the diet (Δaccuracy of 4.5%), presence or absence of a sting (Δaccuracy of 2.3%) and presence or absence of reproductive workers (Δaccuracy of 1.7%) as the most important features for the prediction of trophallaxis behavior (Supplement Fig 1).

With our trained model we predicted trophallaxis behavior for 253 additional species, thus more than doubling the taxonomic coverage for this trait and representing 15 of the 17 ant subfamilies.

### The ancestral ant

To understand the evolutionary context around the emergence of trophallaxis, we used phylogenetic comparative methods and included four additional traits in our analyses for each species: sugary liquids in the diet, workers that can be mated (referred to hereafter as reproductive workers), sting, and colony size. This allowed us to assess the concerted evolution of traits, rates of transition and ancestral phenotypes.

We estimated that the ancestral ant had reproductive workers (posterior probability, pp = 0.983) and a sting (pp = 0.999), while it was unlikely to rely on sugary liquids (pp = 0.037) and did not engage in trophallaxis (pp = 0.002) (Figure 1, Supp Fig 2). The ancestral states of the maximum clade credibility (MCC) tree for colony size were ambiguous, and thus, we cannot assess how the evolutionary history of colony size interplays with the evolution of trophallaxis.

**Figure 1:**
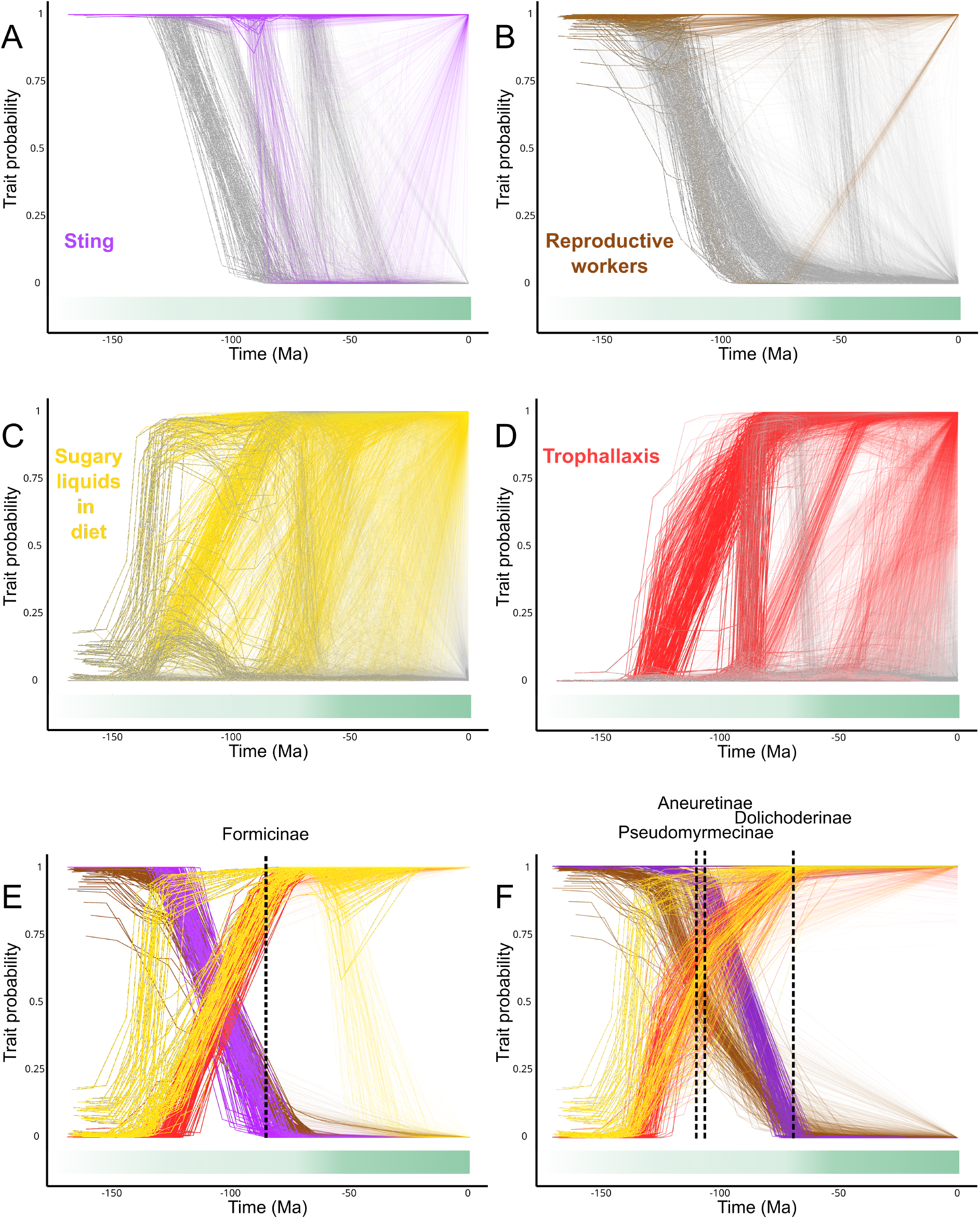
Phenograms showing the probability of presence (p = 1) or absence (p = 0) of phenotypic and behavioral traits in ants inferred through phylogenetic comparative methods. The inferred probabilities are mapped along the branches of the ant phylogeny and plotted against time to show the timing of gains and losses. The traits are A) sting, B) reproductive workers, C) sugary liquids presence in the diet and D) trophallaxis behavior. All traits are shown for two large ant clades that acquire trophallaxis early in their evolution, E) Formicinae and F) Aneuretinae, Pseudomyrmecinae and Dolichoderinae. The phenograms are based on a posterior sample of 100 trees. To account for the uncertainty of predicted presence or absence of trophallaxis across living species we used their posterior probability as inferred by the BNN model rather than its binary predictions (see Methods). On each graph angiosperm prevalence is indicated by a gradient of opacity based on Benton 2022^24^.

Our ancestral state reconstructions indicated that workers with full reproductive potential were lost repeatedly in up to six large events between 125 and 90 Ma, and in up to seven events around 60-50 Ma. Gamergates (a form of fully reproductive workers) were secondarily acquired once in the lineage leading to *Metapone madagascarica* (Myrmicinae) (Fig 1B, Fig 2). Sting was lost at least nine times starting between 115 and 90 Ma and was secondarily evolved at least four times, all within the genus *Stenamma* (Myrmicinae) (Fig 1A).

**Figure 2:**
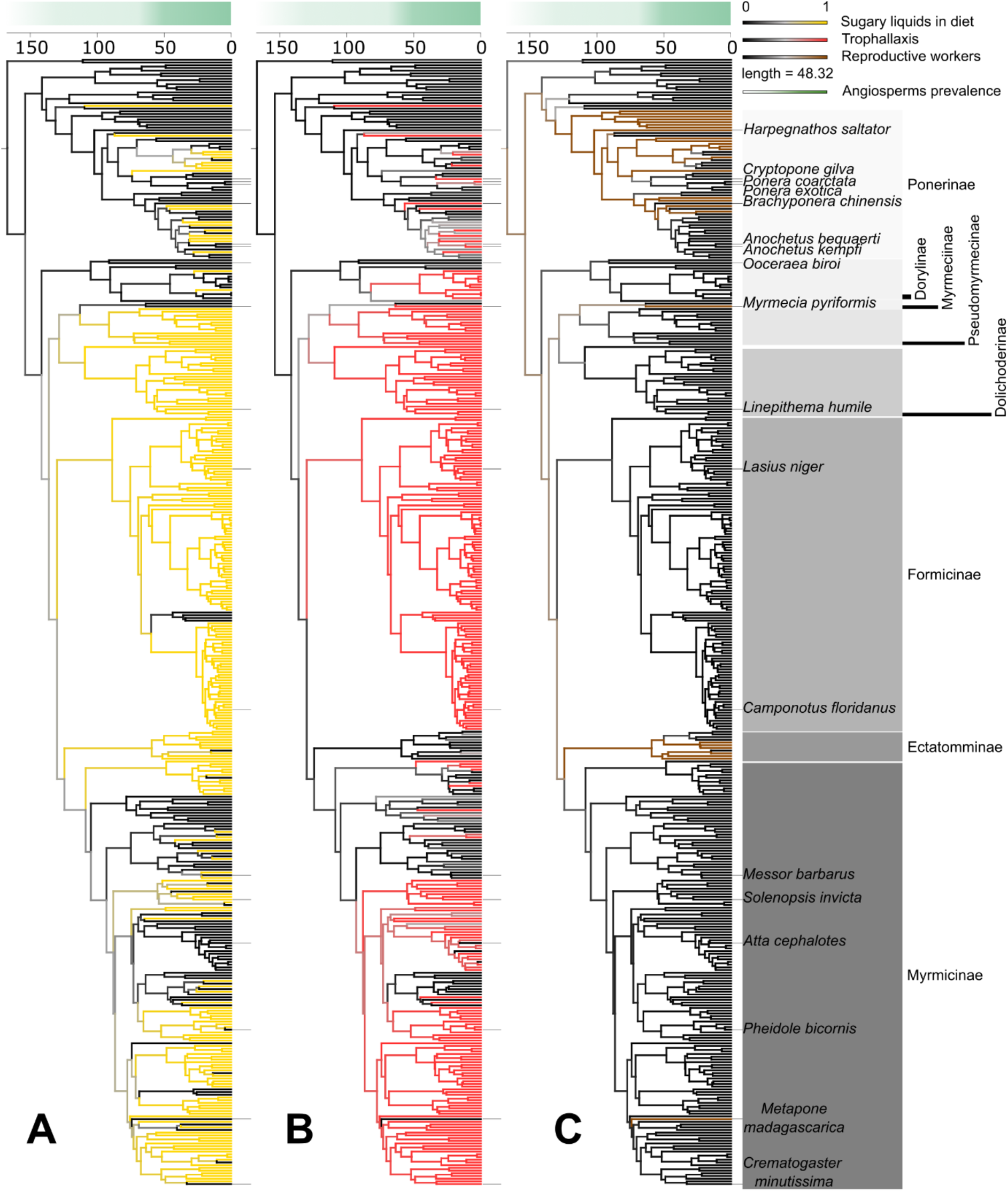
Ancestral state reconstruction integration of A) sugary liquid food in the diet, B) trophallaxis, and C) presence of reproductive workers on the MCC tree for 416 ant species. Colored branches indicate that the trait is likely present at a given time (in Ma) and black that it is absent. Major subfamilies of interest are labeled. The green gradient illustrates the prevalence of angiosperms as described in Benton & al. 2022^24^.

The integration of sugary liquid in the ant diet likely evolved independently multiple times, in few events occurring 140-100 Ma and many more sporadic events 75-10 Ma (Fig 1C, Fig 2). The first event temporally overlapped with a phase of rapid diversification of early flowering plants (~110 Ma)^23^, and the latter with the time when angiosperms became dominant following the End Cretaceous mass extinction (66-50 Ma)^24^. These ecological shifts led to the origination of new biomes and ecosystems for ants to conquer, new food sources for sap-suckers to exploit, and thus highly abundant low-risk food sources (nectar and honeydew) upon which ants could forage.

Mirroring the adoption of sugary liquid in the diet, we found that two of the five biggest ant subfamilies evolved trophallaxis approximately 110 Ma (Forminicae and Dolichoderinae, Fig 1E and 1F, Fig 2). This was followed by more sporadic losses and gains of trophallaxis in other subfamilies throughout the Cenozoic (Fig 1D).

### Conditions for the evolution of trophallaxis

Analyzing the coherent timing across the evolution of traits in different ant lineages, a pattern emerges for the conditions that allowed this social transfer to evolve. The most frequent sequence of evolved traits observed across all our ant species was the early (140-90 Ma) integration of sugary liquid in the diet, the early loss of worker reproductive potential (100-110 Ma) and the early evolution of trophallaxis (120-90 Ma). This sequence of events occurred in the two distinct early evolutions of trophallaxis (Fig 1E and 1F, Fig 2). These two events make up four ant subfamilies (Pseudomyrmeciinae, Dolichoderinae, Aneuretinae and Formicinae) that together account for ~36% of all ant extant species. This leads us to the hypothesis that trophallaxis behavior, a novel social transfer of material^7^, only evolves given the pressure to share liquid food and when intra-colonial reproductive conflict is sufficiently reduced.

Looking across the ant phylogeny, most ant lineages that did and did not evolve trophallaxis did so in accordance with our hypothesis. In the two early gains of trophallaxis (Fig 1E and 1F, Fig 2), ants first began opportunistically eating sugary liquids when these novel food sources became abundant (Fig 2A). Shortly after or at the same time, trophallaxis evolved (Fig 2B), likely as a safe and efficient way to transport and share these new liquid food sources, but trophallaxis only evolved early on in lineages that had also reduced intra-colonial conflict by blocking worker reproductive potential (Fig 2C). The gain of liquid food in the diet alone was insufficient to gain trophallaxis behavior as evidenced by Ectatomminae (Fig 2 & Supp Fig 3). Ectatomminae began drinking sugary liquids during the early diversification of flowering plants, but none evolved trophallaxis. Some Ectatomminae species lost worker reproductive potential rather late, about 40-25 Ma, and yet they still do not perform true trophallaxis. This indicates that there was either some form of critical period connected to ecological changes, or that these species may be transitioning to using trophallaxis, perhaps through mandibular pseudotrophallaxis^16,25^. In multiple other subfamilies, the species present in our dataset never evolved trophallaxis in accordance with our hypothesis (Leptanillinae, Apomyrminae, Proceratiinae, Amblyoponinae and Agroecomyrmecinae, Fig 2): sugary liquids were never integrated into the diet, workers lost full reproductive potential, and trophallaxis never evolved. This indicates that the reduction of intra-colonial conflict is insufficient to drive the evolution of trophallaxis.

In Ponerinae and Myrmicinae, the evolutionary patterns of our four traits are complex, as almost all traits evolved and/or were lost multiple times over a large range of dates. Our ancestral state reconstructions are ambiguous about whether the most recent common ancestor of Myrmicines had already begun eating sugary liquids (Fig 2, Supp Fig 3), and thus conclusions are difficult for this particular subfamily. Ponerine ants rarely perform trophallaxis (16/56 species in our dataset), but those that did evolve trophallaxis also lost worker reproductive potential somewhat consistent with our hypothesis. In Myrmicines, there are many genera and species whose trait evolution is consistent with our model (e.g. *Tetramorium).* However, we also have many Myrmicinae species not known to drink sugary liquids but known or predicted to use trophallaxis, such as *Pheidole bicornis* or *Crematogaster minutissima.* One species, from the subfamily Myrmiciinae, *Myrmecia pyriformis* is a clear exception to our model, in that they have reproductive workers, do not drink sugary liquids and use trophallaxis^26^.

### Traits shaping evolution of trophallaxis

To understand how these traits correlated with trophallaxis over the evolutionary history of ants, we used a phylogenetic d-test^27^ (Fig 3A). We found that diet including sugary liquid (p statistic = 0.002), absence of reproductive workers (p statistic = 0.009), absence of sting (p statistic = 0.016) and large colony size (p statistic = 0.041) all correlated with trophallaxis. Small (p statistic = 0.136) and medium (p statistic = 0.182) colony sizes did not.

**Figure 3:**
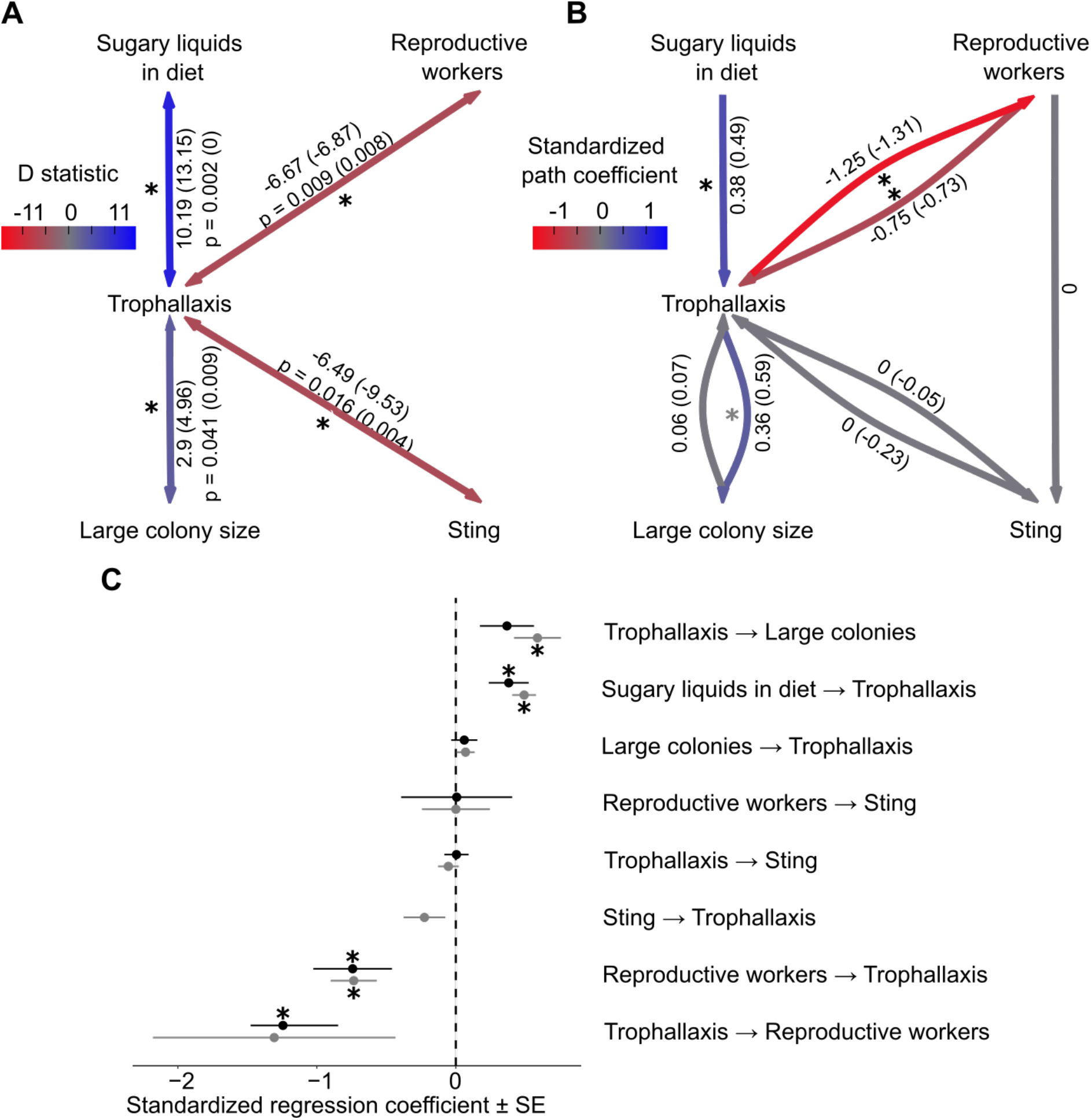
Representation of correlation and causality of traits with trophallaxis. A) Correlation of traits with trophallaxis according to d-test with the D statistic when analyzes are launched on the known and verified 211 species, and on all species including those with imputed data in parentheses and the corresponding p-values. B) Phylogenetic path analysis based on the PGLS model-averaging with the standardized regression coefficient when analyzes are launched on the known and verified 211 species, and on all species including those with imputed data in parentheses. Nodes indicate the presence of the trait. Red edges indicate a negative correlation while blue edges mark a positive correlation. C) The standardized path coefficient between traits and standard error for the phylogenetic path analysis only on the 211 species for which we know the trophallaxis behavior, either from literature or experts (in black) or on the full dataset, including imputed data (in gray). Stars indicate significance based on 95% confidence intervals for one (gray) or both datasets (black).

To disentangle causality in these correlations, we used phylogenetic path analysis using phylogenetic generalized least squares and d-separation^28^. We built models of the trophallaxis-correlated traits based on the results of the d-test (Supp Fig 4 & 5), and the best models, according to AIC, were averaged (Fig 3 B and C). We found that sugary liquids in the ant diet had a significant and positive effect on trophallaxis behavior (effect size 0.38 on only known data, 95% CI [0.09; 0.66]), consistent with our observations given the ancestral states. Trophallaxis, in turn, increased the likelihood of large colony sizes (0.36, 95% CI [−0.02, 0.74]), but large colony sizes did not increase the likelihood of trophallaxis (0.06, 95% CI [−0.13, 0.24]). This indicates that trophallaxis likely did not evolve as a means to better coordinate large colonies, but rather may have improved coordination once the behavior had already been adopted for food sharing. Trophallaxis behavior strongly promoted the loss of reproductive workers (−1.25, 95% CI [−2.02; −0.47]), and reproductive workers strongly inhibited the evolution of trophallaxis (−0.75, 95 % CI [−1.3; −0.19]), indicating that reproductive conflict is likely a much more potent inhibitor of social sharing than previously considered. The results of both the correlation and path analyses on the evolution of sting indicate that while lack of sting may correlate with trophallaxis, there is unlikely to be a causal relationship.

### Speciation and extinction analyses

Given that some of the most species-rich lineages of ants engage in trophallaxis, we tested whether trophallaxis could have impacted speciation and extinction events over the ant phylogeny. Using a Bayesian Analysis of Macroevolutionary Mixtures (BAMM)^29^ with STructured Rate Permutations on Phylogenies test (STRAPP)^30^ we found that trophallaxis significantly lead to speciation events (p = 0.0423), with lineages engaging in trophallaxis showing 1.6x higher speciation rates (no trophallaxis = 0.07±0.004; trophallaxis = 0.11±0.007) and 2.8x higher extinction rates (no trophallaxis = 0.007±0.004; trophallaxis = 0.02±0.007). However, when accounting for the possibility of other, unmeasured traits affecting diversification using a state-dependent model with hidden states (HiSSE)^31^, we found that the preferred model implied character independent diversification with three hidden states (delta AIC > 8000; Supp Table 3). Thus, the HiSSE model rejects a direct link between trophallaxis and increased speciation rates. While this discrepancy between methods can derive from model identifiability issues ^32,33^, it can also indicate that trophallaxis might have played an indirect role in ant diversification or that speciation patterns cannot be ascribed solely to this trait.

## Discussion

Our results suggest that trophallaxis behavior evolved to allow transport of sugary liquids that became abundant first during the diversification of angiosperms and more so later when angiosperms became dominant. This represents a key shift from a specialist diet (on insect prey) to a generalist diet (including both prey and sugary liquids). This type of dietary shift, in other systems, has been linked to faster diversification^34,35^, range expansion^36,36,37^ and invasion^38–41^. In spite of clear advantages to becoming a generalist, not all ants did so in their early evolutionary history when sugary liquids became abundant – ant lineages that retained the worker ability to reproduce did not evolve the social transfer of trophallaxis. Only the ant lineages that became sugar-eating generalists and also reduced intra-colonial conflict early in their evolutionary history, losing the worker capacity to reproduce, evolved trophallaxis. Extrapolating from our dataset to the extant number of species in these clades, this sequence of trait evolution likely occurred in the evolution of about 36% of all ant species, the most consistent pattern of trait evolution over time across our dataset.

Behavior has been proposed to both drive evolution and to buffer it^1^ and indeed there is evidence for both in different contexts. Studying the impact of behavior on macroevolution is often difficult because we cannot easily measure behavior from fossils and it can be difficult to access some extant species. Here we have adapted new deep-learning tools to impute behavior, allowing us to bridge this gap and study the evolution of a complex behavior over a large phylogeny. Bayesian neural networks provided a flexible tool to integrate multiple traits along with phylogenetic structure in our predictions, while embracing and explicitly quantifying the uncertainties around them. In our analytical framework these uncertainties were then propagated to the comparative phylogenetic analysis and ancestral state estimations to provide robust inferences.

Trophallaxis likely first evolved in ants as a mode of sharing exogenous food, yet in ants that perform this behavior frequently (e.g. *Camponotus*), trophallactic fluid has become a rich and complex socially transferred material similar to mammalian milk^17,19^. Various proteins and small molecules used elsewhere in ant physiology have been co-opted and neo- or sub-functionalized as they became part of trophallactic fluid^18^. Juvenile hormone and juvenile hormone esterase provide an example of developmental hormones and hormone regulators that are critical in insect development, that once co-opted to trophallactic fluid, allow workers to manipulate larval development through adult-to-larva trophallaxis^17,18^. Once a social transfer, like milk-feeding or trophallaxis, evolves it creates a private physiological channel between bodies that has far reaching implications for social life and the evolutionary and ecological success of social organisms^7^. A potentially important consequence of the evolution of trophallaxis is the possibility of an individually tailored diet for larvae, which more easily allows body-size variation between otherwise genetically equivalent individuals. Such asymmetries are hallmarks of committed cooperative systems after major evolutionary transitions in individuality^42^, as they can entrench a cooperative system into rigid specialization.

Our scenario for the evolution of trophallaxis in ants differs from the evolutionary path inferred for wasps, another eusocial hymenopteran lineage^43,44^. In wasps, trophallaxis likely evolved from adults pre-chewing prey and providing it to larval young, and generally, adult-adult trophallaxis only occurs in highly eusocial wasps^43^. In ants, because the data on larval-adult trophallaxis are so sparse^14^, we were not able to explore the possibility that another social transfer may have played a role in the evolution of trophallaxis. Adult-adult stomodeal trophallaxis alone is truly a continuous trait rather than a binary one, where some species use the behavior only in the context of food (most Ponerines that engage in trophallaxis), while others perform it routinely across contexts (*Camponotus).* Looking at our model’s predictions for the Myrmicinae or Dorylinae (our BNN predicted that New World army ants use trophallaxis, even though, to our knowledge, trophallaxis behavior has never been reported), it is possible that the use of some other form of social transfer has led to our model’s uncertainty or unexpected predictions. Further investigation of these clades would help verifying the predictions and refining the model.

Our result, that ancestral ants likely had workers that could mate, brings significant doubt to the idea that all ants are superorganismal. It has previously been postulated that all ants are superorganismal^13^ because they have passed a ‘point-of-no-return’ of having morphologically fixed reproductive division of labor. A main point of evidence for this comes from fossils of ants from a sister clade to crown Formicidae where winged queens and unwinged workers were fossilized together in the same group^45^. However, the winged individuals may have been dispersal morphs while workers may have retained reproductive potential in the form of a functional spermatheca^46^. These two conditions are met in the extant ant *Harpegnathos saltator* known to have both “true queen” (winged) and gamergates^47^ (reproductive workers). Others have also noted this potential ambiguity in the fossil record^48^. Investigations of fossil worker spermathecae could indicate whether ancestral ant workers could be mated. Our result suggests that gamergates are not secondarily evolved^49^, but rather that reproductive workers were present in the common ancestor to all ants. This, in turn, means that according to Boomsma’s definition of superorganismality, the common ancestor to all ants had not passed this point of no return and that fully reproductive workers may have been retained as a plesiomorphic trait, and thus not all ants are superorganismal. Our results indicate instead that superorganismality evolved multiple times within Formicidae, presenting a rich playground to study major evolutionary transitions in individuality^42^.

Our study demonstrates a link between environmental changes brought about by the rise of flowering plants, with its transformational impact on most terrestrial ecosystems, and a simple but important ant behavior that had notable impact on the evolution of different ant lineages. This shift was enabled by ant-plant and ant-sap-sucker interactions. These resulted in adaptations across multiple partners, and thus significant rewiring of ecosystems^3,50^. Ants, and in particular ants that rely on such mutualisms, are known to be ecosystem engineers, moving their aphid cattle from plant to plant, protecting trees from predators, cultivating fungus gardens and planting seeds. Understanding the evolutionary underpinnings of behavioral traits is crucial to grasp their fundamental role in determining the ecological success of the most dominant organisms in the animal kingdom, such as *Homo sapiens* and the ants.

## Methods

### Data collection

We completed the reports of trophallaxis from Meurville & LeBoeuf^14^ by manually searching literature up to early 2021 for reports of occurrence or absence of trophallaxis behavior, ending up with data for trophallaxis behavior of 205 species from 265 sources (Supp Table 1). Because some subfamilies were more represented than others, and in order to have more even taxon sampling, we imputed trophallaxis behavior (Supp Fig 6). To do so, we used a Bayesian Neural Network (BNN) as implemented in the npBNN library^51^ (https://github.com/dsilvestro/npBNN), that provided posterior probabilities for a binary trait, based on life-history, morphological and ecological traits collected from literature. These traits were either easy to collect for a large number of species (temperature and humidity data, presence or absence of a sting) or were linked to some of our hypotheses (Supp Table 4).

### Phylogeny

We used the MCC tree produced by Economo et al. 2018^52^ as a reference for the ant phylogeny of over 14,000 species, as it was the largest and most recent ant phylogeny at the time. We corrected species names according to antwiki (https://www.antwiki.org/wiki/Ant_Names) valid species names in 2021, and removed morphospecies. In order to include the phylogenetic signal in the BNN to predict trophallaxis behavior, we transposed the tree into eigenvalues using the R packages ape v5.4.1 and PVR v0.3, providing us with 11 eigenvalues (50% of the phylogenetic signal) per species that we used as an input to the BNN.

### Temperature and humidity

We retrieved temperature and humidity data from the GABI database (Release 18.01.2020) and Antweb (October 2020). Temperature and precipitation data were obtained from the coordinates referenced in the two aforementioned databases, using the R package raster v3.3.13. For each specimen of each species, we collected the annual temperature and precipitation recorded at this location. For each species, we then extracted the minimal and maximal values of temperatures and precipitations, thus providing a range of temperatures and precipitations for each species’ habitat, ending in four continuous traits: minimal and maximal temperature and minimal and maximal precipitation.

### Drinking sugary liquids

Mention of a species drinking sugary liquids in the field, tending aphids, or having repletes was reported as 1, and specialized predators or ants not interested in sugary foods were attributed a 0. For four species we accepted information indicating species drink sugary liquids in the lab (*Strumigenys lewisi, S. canina*, *Cephalotes varians* and *Streblognathus aethiopicus*). For 32 species in 17 genera (*Acanthognathus, Anochetus, Apomyrma, Cerapachys, Cryptopone, Discothyrea, Leptogenys, Myopias, Myrmecina, Neivamyrmex, Platythyrea, Ponera, Prionopelta, Probolomyrmex, Proceratium, Typhlomyrmex* and *Tapinoma*), the species-level diet information was impossible to retrieve, so it was inferred from the genus-level.

### Sting

Whether species have a sting or not was retrieved from the supplementary material of Blanchard & Moreau^53^. As sting is not considered to be an evolutionarily labile trait, we extrapolated the data on presence or absence of a sting from genus to species level. Missing values in the training set were retrieved manually from scientific literature (Supp Table 4).

### Colony size

Colony size data were retrieved manually from various sources. Because colony size is variably reported in the literature (sometimes maximum, minimum, average or single observations), we recorded what was available. To face this heterogeneity in data collection, we split colony size by quartile (1st quartile: under 60 individuals, 2nd quartile: 60-268, 3rd quartile: 269-2000, 4th quartile: 2001-306000000). To binarize the data, each quartile was considered a trait.

### Reproductive workers

Categorical data on presence or absence of gamergates (ants with worker fate but that can be mated and produce diploid offspring) was retrieved from antwiki, https://www.antwiki.org/wiki/Category:Gamergate (October 2021). Genera not listed were assumed to have workers that could not be mated. Species from genera known to have gamergates but for which no data was found were not included in the dataset, as we do not know how labile the presence or absence of gamergates is within a genus.

### Data imputation

We trained a BNN model on 161 species split as training (90%) and test sets (10%). To evaluate the test accuracy of the model we performed a 10-fold cross-validation analysis. The BNN model included a fully connected network with two hidden layers of 20 and 10 nodes, respectively, with ReLU activation function^54^.

The cross-validation test accuracy of our model was 84% (Supp Table 2). We had, on average over the 10 test-samples, 11.6 True Positives, 1.8 True Negatives, 1.6 False Positives and 1 False Negative.

To further evaluate the accuracy of our model, we sent the list of species for which we predicted trophallaxis behavior (including both predictions that species would engage in trophallaxis and predictions that a species would not engage in trophallaxis) to 10 expert myrmecologists (C. Lebas, N. Idogawa, A. Yamada, P. Slingsby, R. Mizuno, J. Heinze, A. Lenoir, V. Nehring, F. Savarit and X. Cerda, Supp Table 5) and asked them if they had observed or not observed stomodeal trophallaxis between adults for any species on the list. We used their answers to estimate the accuracy of our imputations. Experts provided responses for 48 species, 19% of our imputed species. We have 37 True Positives, 5 True Negatives, 3 False Positives and 3 False Negatives, leading to a myrmecologist accuracy of 88% (Supp Table 2).

After running the trained model on 253 unlabeled species, we obtained a dataset of 416 species: 161 species with known trophallaxis behavior and all known traits, 2 species with known trophallaxis behavior but missing temperature and precipitation data (*Myrmicaria natalensis eumenoides* and *Myrmoteras jaitrongi,* Supp Table 4), 48 species with trophallaxis behavior confirmed or denied by myrmecologists, and 205 species with trophallaxis behavior imputed by the BNN. In cases of mismatch between our predictions and the experts’ assessments, we gave priority to the trophallaxis behavior indicated by the experts.

### Evolutionary analyses

We used sMap v1.0.7^55^ to infer ancestral states using phylogenetic comparative methods. For each trait (presence or absence of reproductive workers, sting, trophallaxis and sugary liquid food in species diet; discretized colony size) we tested different Markov models in a maximum likelihood framework and ranked them based on Akaike weights (Supp Fig 7).

Analyses were launched with 1000 simulations using the ‘sMap’ command. We launched maximum likelihood analyses on both the stem maximum clade credibility tree and 100 stem posterior trees in order to take variability into account (Supp table 6). The rest of the analysis was done only on the maximum clade credibility tree. We then built a Bayesian model with a log normal prior where μ is the natural logarithm of the maximum likelihood model rates and σ is 1. We then computed the posterior probability of each model and used it to average Bayesian models with the ‘Blend-sMap’ command. This provided the ancestral state reconstruction for the stem maximum clade credibility tree.

### Trait correlation and phylogenetic path analysis

We used a d-test^27^ as implemented in sMap^55^ to estimate the co-occurrence of each trait with trophallaxis. We performed this test on the known dataset and verified species (211 species) to avoid circularity and on the full dataset (416 species). We performed the ML analysis and Bayesian analyses as before, only adding the -pp 100 option to draw 100 posterior predictive samples. Then, we average Bayesian models and pairwise merged the trophallaxis model with every trait using the ‘Merge-sMap’ command to finally use ‘Stat-sMap’ with default values to run the D-test.

To test how traits influenced trophallaxis behavior, we used phylogenetic path analysis as proposed by von Hardenberg and Gonzalez-Voyer^28^, on our dataset of 416 species, and on only the 211 species with known trophallaxis behavior (from literature and imputations verified by myrmecologists). This method allows the comparison of possible causal relationships between traits while testing for direct or indirect effects and considering the non-independence of the traits due to phylogeny. We first built causal quantitative models as directed acyclic graphs (DAGs, Supp Fig 4) and accounted for phylogeny through phylogenetic generalized least squares analysis (PGLS), using the d-separation method^28,56^. Building the models, running the phylogenetic path analysis and performing model selection was done in R with the Phylopath v.1.1.3^57^, in order to take both continuous and discrete response variables into account, with 500 bootstrap.

The best models were models seven, one, three and five for the full dataset, and models one, three and seven for the reduced dataset (Supp Fig 4 and 5). Because we cannot identify the best model, we averaged the models using the average() function in the phylopath package. This provided a model that reflects the results of the best models.

### Speciation and extinction analyses

We used reconstructed birth-death processes to estimate the dynamics of speciation and extinction rates across the ant phylogeny and assess whether the evolution of trophallaxis might have affected species diversification. To this end we used the state-dependent models implemented in the R package HiSSE (v 1.9.6)^31^ and the mixture models implemented in BAMM (v 2.5.0)^58^.

For the HiSSE analysis, we used the tree from Economo et al 2018^52^ tree with 14,183 species, and set the trophallaxis state to unknown for all species missing from our imputed dataset. We tested four models (Supp table 3), starting with (1) the original BiSSE model^59^, with turnover (speciation + extinction) and extinction fraction (extinction / speciation) dependent on that trait state. Because the BiSSE model is prone to false positives^31^, we also ran a character-dependent HiSSE model (2) with two hidden states, where diversification rate variation can occur as function of both trophallaxis and an unobserved (hidden) trait. We additionally tested character independent models with only hidden traits (of two or three) affecting turnover and extinction fraction and with no effect of trophallaxis. We compared the fit of these models through maximum likelihood using AIC.

For the BAMM analysis, we started estimating empirical priors using the setBAMMpriors function from BAMMtools v2.1.9, and launched BAMM on a reduced phylogeny for the 416 species in our dataset with species-specific sampling fractions determined at the genus level and default parameters for other arguments (Supp Table 7). We then performed a STRAPP analysis^30^ test with the BAMMtools function traitDependentBAMM, to estimate if rates differ between species using and not using trophallaxis. We used the resulting P-value to assess whether significant evidence of a trait-dependent effect was detected and the mean diversification rates across species engaging or not with trophallaxis to quantify the magnitude of that difference.

## Acknowledgments

We thank Corrie Moreau and Robert Waterhouse for their precious insights on this project, Giorgio Bianchini for his support in using sMap, and Torsten Hauffe for his support on the speciation/extinction analyses. We also thank the expert myrmecologists Claude Lebas, Naoto Idogawa, Aiki Yamada, Peter Slingsby, Riou Mizuno, Juergen Heinze, Alain Lenoir, Volker Nehring, Fabrice Savarit and Xim Cerda who kindly took the time to share their knowledge about trophallaxis with us.

## Author contributions

M-P.M, D.S and A.C.L conceived the experiments. M-P.M and A.C.L collected the traits data from literature. M-P.M conducted the analyses under the supervision of D.S and A.C.L. M-P.M, D.S and A.C.L made the figures and wrote the manuscript with comments from all authors.

